# Data mining redefines the timeline and geographic spread of cotton leafroll dwarf virus

**DOI:** 10.1101/2024.06.05.597610

**Authors:** Alejandro Olmedo-Velarde, Hayk Shakhzadyan, Michael Rethwisch, Michael West-Ortiz, Philip Waisen, Michelle Heck

## Abstract

Cotton leafroll dwarf virus (CLRDV), a threat to the cotton industry, was first reported in the United States (US) as an emergent pathogen in 2017. Phylogenetic analysis supports the hypothesis that US CLRDV strains are genetically distinct from strains in South America and elsewhere, which is not consistent with the hypothesis that the virus is newly introduced into the country. Therefore, using database mining, we evaluated the timeline and geographic distribution of CLRDV in the country. We uncovered evidence that shows that CLRDV had been in the US for over a decade before its official first report. CLRDV sequences were detected in datasets derived from Mississippi in 2006, Louisiana in 2015, and California in 2018. Additionally, through field surveys of upland cotton in 2023, we confirmed that CLRDV is present in California, which had no prior reports of the virus. Viral sequences from these old and new datasets exhibited high nucleotide identities (>98%) with recently characterized US isolates, and phylogenetic analyses with their homologs placed these sequences within a US-specific clade, further supporting the earlier presence of CLRDV in the country. Moreover, potential new hosts, including another fiber crop, flax, were determined through data mining. Retrospective analysis suggests CLRDV presence in the US since at least 2006 (Mississippi). These findings necessitate a reevaluation of spread patterns, inoculum sources, symptomology variations, and control strategies. Our findings challenge the current understanding of the arrival and spread of CLRDV in the US, highlight the power of data mining for virus discovery, and underscore the need for further investigation into CLRDV’s impact on US cotton.

## Main Text

Cotton leafroll dwarf virus (CLRDV) is a member of the genus *Polerovirus* (Family *Solemoviridae*) that represents a persistent threat to the United States (US) cotton industry (Edula et al. 2023). Since its first reported detection in Alabama in 2017 (Avelar et al. 2019), it has spread to almost all the states in the cotton belt except for California, Arizona, New Mexico, and Missouri (Brown et al. 2019; Edula et al. 2023). CLRDV can cause significant yield losses in several, but not all, states in the cotton belt region where its presence has been confirmed (Brown et al. 2019; Mahas et al. 2022).

The symptoms associated with the virus are complex and vary depending on biotic and abiotic factors, such as cultivar, environment, and stage of growth, and can be aggravated by other underlying stress (Parkash et al. 2021). Nevertheless, symptoms may include interveinal chlorosis, leaf rolling and distortion, stunting, bronzing or reddening, and reduced boll sets (Edula et al. 2023). Distinct strains of CLRDV have been regarded as the causal agent of cotton leafroll dwarf virus disease (CLRDD) in the US (Brown et al. 2019) and Thailand (Sharman et al. 2015) and cotton blue disease (CBD), previously known as cotton vein mosaic disease (Ferguson and Ali 2023), in South America (Distefano, Bonacic Kresic, and Hopp 2010; Agrofoglio et al. 2017). CLRDV may have originated in Africa and may underlie CBD on the continent (Ferguson and Ali 2023). This hypothesis previously originated since what is called CBD in Africa is characterized by very similar symptoms to those reported in South America (Cauquil 1977) and is spread by aphids (Cauquil and Vaissayre 1971). However, neither serological nor molecular studies have been performed on cotton presenting CBD symptoms in Africa to confirm that CLRDV causes CBD (Ferguson and Ali 2023).

Therefore, the question remains unanswered regarding whether CLRDV is an exotic virus recently introduced to the US or if its presence in cotton production went unnoticed until the emergence of a new strain within the country, influenced by several factors such as frequent polerovirus recombination (LaTourrette, Holste, and Garcia-Ruiz 2021). Data mining of publicly available datasets has emerged as a powerful approach to virus characterization and discovery, leveraging our knowledge of plant virus diversity (Bejerman, Dietzgen, and Debat 2021; Bejerman, Dietzgen, and Debat 2023; Edgar et al. 2022) and their geographic locations (Rivarez et al. 2023; Edgar et al. 2022). Here, we used two data mining approaches to retrieve CLRDV sequences from archived datasets in publicly available databases in GenBank. We report evidence consistent with CLRDV’s presence in the US for more than ten years before its official recognition. We also expand its geographic location to a new host and a new state in the cotton belt, validated using serological and molecular diagnostic approaches, and also add potential new natural hosts of the virus. This revised timeline holds significant implications for understanding the virus’s spread, potential sources of inoculum, symptomology caused by the virus, and the effectiveness of current control strategies.

In the first data mining approach, we explored the Serratus database through the Serratus Explorer and the palmID Viral-RdRp analysis tools. Serratus is an invaluable tool for virologists. It acts as a powerful data mining platform, allowing users to explore a vast collection of viral sequences. Specifically, Serratus focused on Sequence Read Archive (SRA) accessions up to January 2020 that contain predicted viral RNA-dependent RNA polymerase (RdRp) motifs (Edgar et al. 2022). For both tools, we used the CLRDV P1-P2 protein (YP_003915148) as a query with a score alignment >50 and protein identities >90%. Using Serratus, we retrieved a total of 114 unique SRA libraries whose assembled reads (contigs) potentially had CLRDV RdRp motifs. A manual examination of the STAT-generated analysis (Katz et al. 2021) of each SRA library accessed through the SRA Run Selector website found 0.000005% (135 reads) through 0.693% (394,555 reads) of the total number of reads, which had CLRDV origin. This corroborates the presence of the virus in all the mined datasets (Table S1). Most of these SRA libraries originated from China from *Gossypium hirsutum, G. barbadense*, a *Gossypium* hybrid (*G. hirsutum* x *G. australe*), and *Hibiscus cannabinus* (Figure 1A and Table S1). Even though CLRDV has been previously reported naturally infecting other malvaceous hosts in Asia, such as *Hibiscus syriacus* (Igori et al. 2022) and *Malvaviscus arboreus* (Wang et al. 2023), we identified *Hibiscus cannabinus* as a possible new natural host for CLRDV. CLRDV was initially reported in soybean aphids collected from China in 2009, even though cotton is a non-preferred host of this aphid species (Feng et al. 2017). Subsequently, the virus was officially reported in mixed infections with the aphid-borne potyvirus watermelon mosaic virus in cotton plants presenting leafroll and vein yellowing collected in May 2016 (Yang et al. 2021). Interestingly, the earliest sample collection date we found in the datasets from China was July 2015 (Table S1), suggesting CLRDV might have been present in cotton in China before the official report in 2016. The datasets that originated from India were derived from *G. hirsutum, Cicer arietinum* (chickpea), and *Linum usitatissimum* (flax) (Figure 1A). CLRDV was first reported infecting upland cotton in India in plants presenting stunting, leaf rolling, vein yellowing, and reduced boll size or sterility (Mukherjee et al. 2012), and chickpea plants affected with chickpea stunt disease (Mukherjee, Mukherjee, and Kranthi 2016). Therefore, this study adds flax as a new, unexpected fiber crop as a possible host of CLRDV. The datasets from the US were derived from two *G. hirsutum* samples and, unexpectedly, from a bovine gut ‘metagenome’ sample generated from California. One of the *G. hirsutum* datasets corresponds to the first complete CLRDV genome obtained from Texas (Ali and Mokhtari 2020); however, the second dataset corresponds to a transcriptomics study performed in Louisiana in 2015 that aimed to identify genes responding to the seed infection by *Aspergillus flavus* (Bedre et al. 2015). To further evaluate the CLRDV presence in the cotton seed and bovine gut datasets, we retrieved both SRA datasets (SRR1805340 and SRR9035366) and used Trimmomatic 0.38 (Bolger, Lohse, and Usadel 2014) with default parameters to obtain quality-trimmed reads. Reference-guided assembly was performed using the Geneious mapper (Kearse et al. 2012), quality-trimmed reads, and a selected CLRDV isolate from Texas (OK185944). Almost complete CLRDV genomes from the cotton seed (5,823 nt, accession BK068117) and the bovine gut (5,809 nt, accession BK068117) datasets were obtained with mean coverages of 62X and 5X, respectively, throughout almost the entire genomic sequence (Figure 1B). BLAST searches revealed that the almost complete CLRDV genome retrieved from the cotton seed dataset generated from Louisiana in 2015 (LA2015) presented 98.3% nucleotide identity with a Texas isolate (TXb, OK185944), while the one retrieved from the bovine gut dataset generated from California in 2018 (CA2018) presented 97.4% nucleotide identity with a different Texas isolate (CS4, MN872302). Evolutionary relationships of the almost complete genomes of LA2015 and CA2018, and their CLRDV homologs available as of March 2024 were obtained using Bayesian inference. The phylogenetic tree was constructed using MrBayes 3.2.7 (Huelsenbeck and Ronquist 2001) implemented in NGphylogeny.fr (Lemoine et al. 2019). The best evolution model (K2P + G) with three Markov chain Monte Carlo runs of 100,000,000 generations sampling every 10,000 trees was used to infer the phylogenetic relationships. Sequences were aligned using MUSCLE 3.8.1551 (Edgar 2004), and the ambiguous positions in the alignments were curated using Gblocks 0.91.1 (Talavera and Castresana 2007). The consensus tree was visualized and manually edited in iToL 6.8.2 (Letunic and Bork 2021). The Bayesian inference revealed geographically distinct clades, with isolates from the US, Asia, and South America clustering together. Previous studies have found similar topologies in the phylogenetic trees constructed using either partial or almost complete genome sequences (Ramos-Sobrinho et al. 2021; Tabassum et al. 2021). Interestingly, the LA2015 and CA2018 isolates grouped in a clade containing isolates from Texas only, while also being within a larger clade containing isolates from South Carolina, Texas, Alabama, and Florida (Figure 1C).

**Figure 1.**
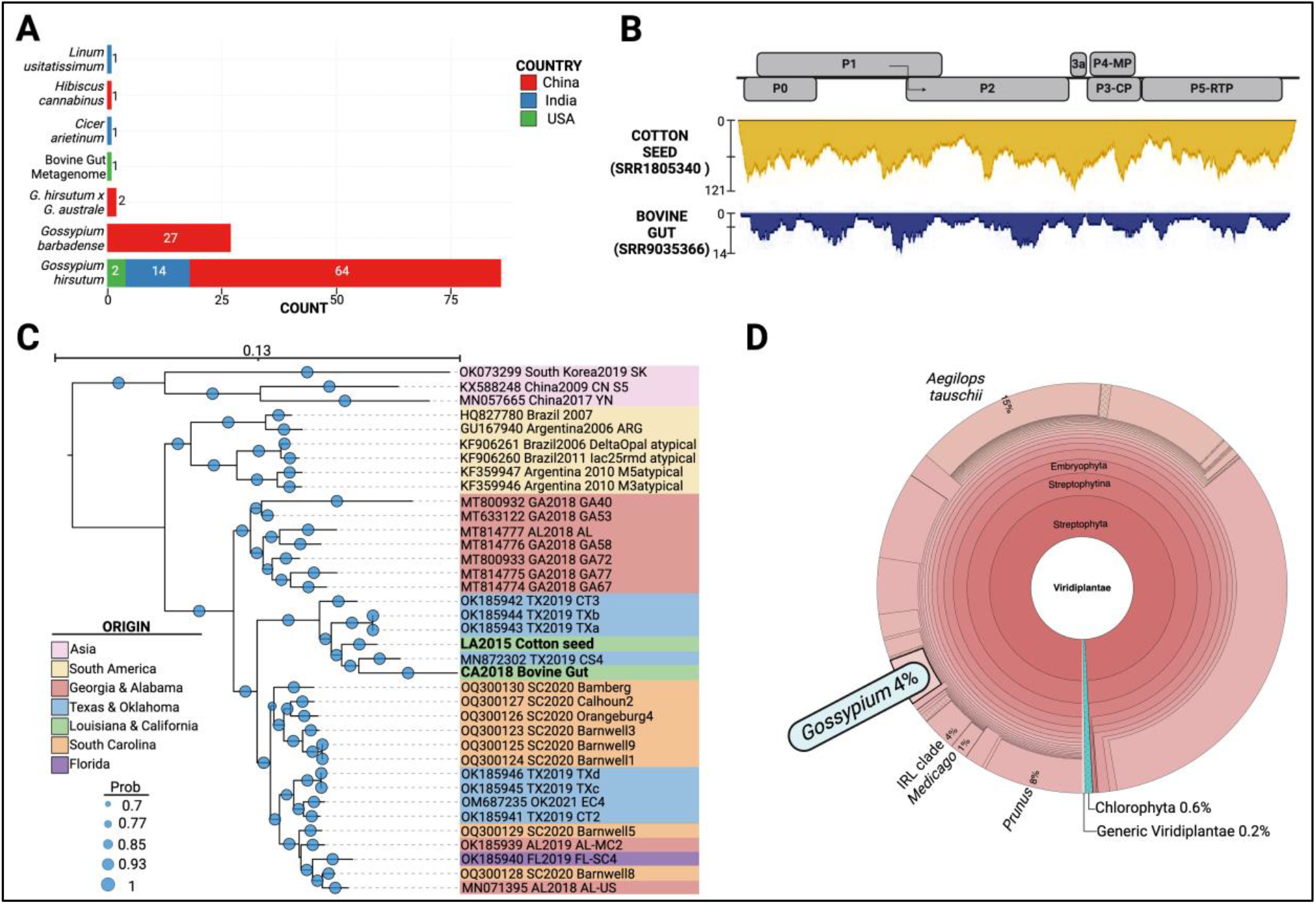
Data mining of cotton leafroll dwarf virus (CLRDV) sequences retrieved from the Serratus (serratus.io) database. A) Count of Sequence Read Archive (SRA) datasets originated from different organisms and countries that have CLRDV RNA-dependent RNA polymerase (RdRp) hallmarks. B) Coverage of quality-trimmed reads mapped against a reference CLRDV genome (OK185944) from two selected datasets: Cotton seed from Louisiana 2015 (LA2015) and bovine gut metagenome from California 2018 (CA2018). C) Phylogenetic relationships of the CLRDV genomes newly assembled from the two SRA datasets from LA2015 and CA2018 (bold). The tree was constructed through Bayesian inference with Mr. Bayes, and the best model of DNA evolution (K2P + G + I). D) Taxonomic analyses of the CA2018 dataset with reads identified with plant (Viridiplantae) origin. About 4% of the plant reads were assigned to *Gossypium*.

CLRDV not only naturally infects the hosts mentioned above but also cacao (Ramos-Sobrinho et al. 2022) and a variety of weed hosts (Sedhain et al. 2021). Therefore, to infer the possible host that was likely infected by CLRDV and fed to the cow for the bovine gut dataset, the STAT-generated analysis for the SRA accession was manually inspected, as detailed above. About 4% of the plant (Viridiplantae) reads were predicted to originate from *Gossypium* (Figure 1D). Since we could not be certain that the cow was not transported from elsewhere to California, where the dataset was generated, or the origin of the *Gossypium* plants fed to the cow, we decided to further evaluate the presence of CLRDV in California, where the presence of the virus has not been confirmed. In September 2023, we visited several upland cotton fields from two locations in southern California: Palo Verde Valley and Coachella Valley (Figure 3A-B). We noted a range of virus-like symptoms in some plants mostly obvious on the borders of the fields. The symptoms were characterized by leaf rolling and crinkling, interveinal chlorosis, taller than normal plants with stacked nodes in the upper canopy, and mosaic (Figure 2C-F). All the symptoms mentioned above have been previously associated with CLRDV (Brown et al. 2019; Edula et al. 2023), except for mosaic. A total of 34 leaf samples with their petiole attached were collected from symptomatic plants, submerged in RNAlater®, and transported to our laboratory at Cornell. The samples were used to initially evaluate the presence of CLRDV by double-antibody sandwich (DAS)-ELISA using camelid single-chain antibodies raised against the coat protein of CLRDV as the capture antibody (Patent USPTO 18/436,287) and the commercially available Anti-PLRV conjugate (Agdia, Elkhart, IN) as the secondary antibody. DAS-ELISA detected CLRDV in a total of 15 symptomatic samples (Table 1). Positive and negative controls tested positive and negative, and consisted of tissue from a CLRDV-infected tree from Mississippi maintained in our greenhouse since 2019 and from healthy upland cotton seedlings, respectively. The presence of CLRDV in these samples was further evaluated by RT-PCR assays. Total RNA was extracted from midrib and petiole samples using the Spectrum Plant Total RNA kit (Sigma), and complementary DNA (cDNA) was synthesized using the iScript Reverse Transcriptase kit, as per the manufacturers’ instructions. A single-tube nested RT-PCR assay implemented to amplify a partial region of the RdRp gene of CLRDV (West-Ortiz et al. Phytobiomes submitted) was used to index the presence of CLRDV in the cDNA samples. The expected 350 bp product was obtained from only five samples (Table 1). Direct Sanger sequencing of three purified PCR products and pairwise comparisons demonstrated they presented 100% nucleotide identities among one another. BLAST searches using the consensus sequence (accession PP806565) showed <98.9% nucleotide identities with isolates from Alabama (MT814777), Georgia (MT633122.1), Texas (OK185943-4), and Argentina (KF359946-7). Considering a sample positive for CLRDV when it was positive using either DAS-ELISA or RT-PCR assays, a total of 17 out of the 34 samples tested were positive for the virus. Most of the symptomatic samples that tested positive for CLRDV commonly presented mosaic symptoms or a group of the symptoms mentioned above (Table 1). Interestingly, the mosaic symptoms observed in the California samples (Figure 2E), possibly caused by CLRDV, highly resemble those caused by cotton bunchy top virus, the closest *Polerovirus* member that infects cotton in Australia (Ellis et al. 2013; Reddall et al. 2004). Collectively, this study provides *in silico* evidence and laboratory confirmation that CLRDV is present in California. Nevertheless, it is necessary to evaluate if the virus is causing any yield affection and if it is also present in the San Joaquin Valley, where most of the production in the state is located.

**Table 1.**
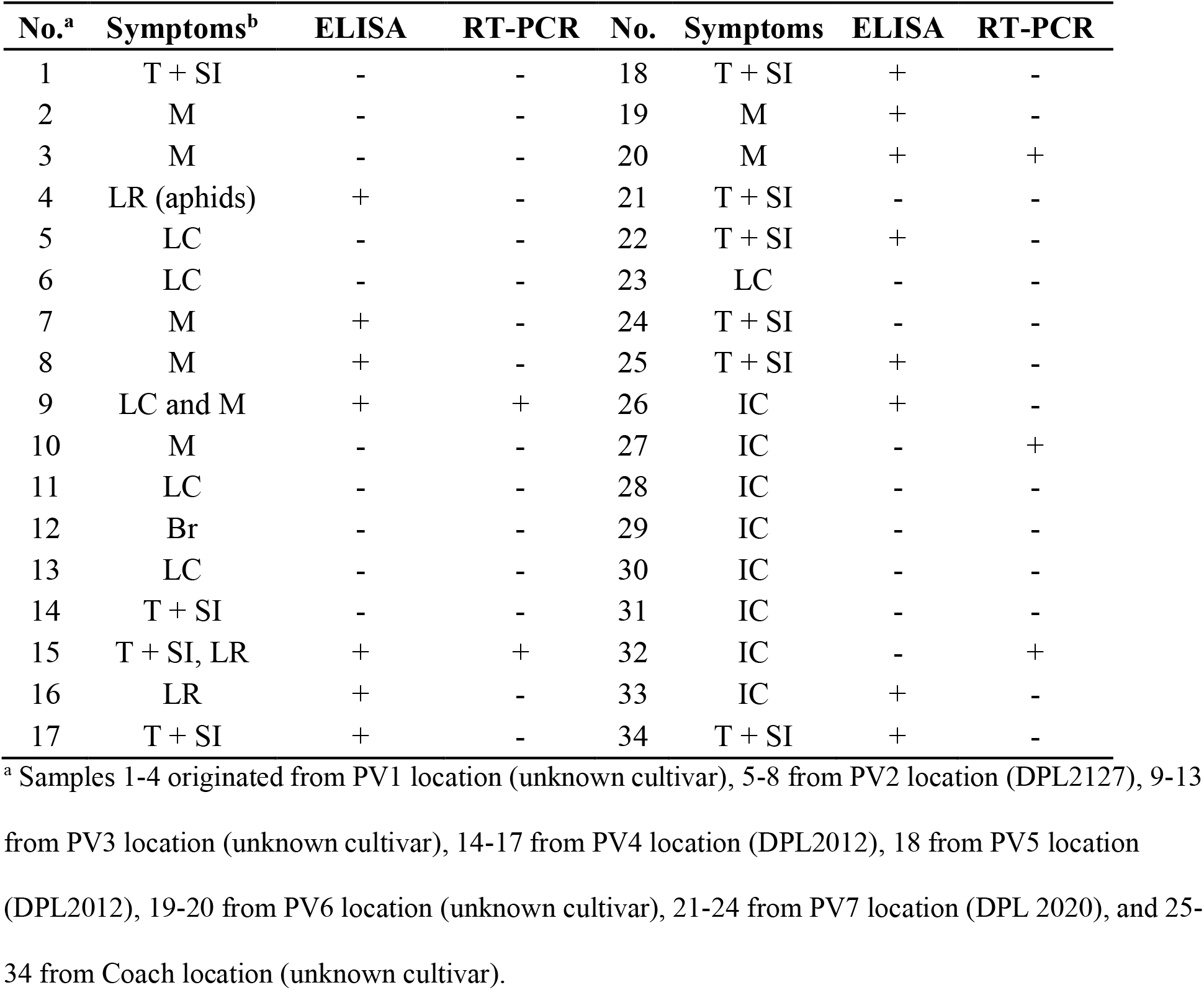
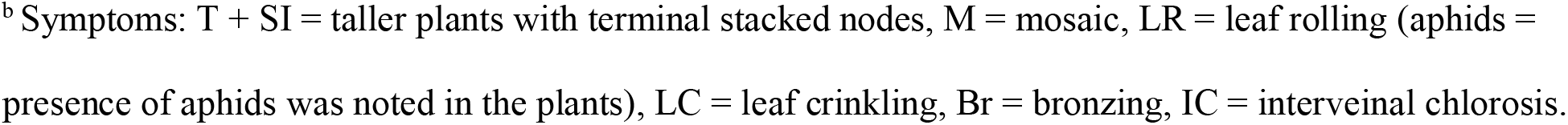
Viral-like symptoms and virus indexing results on 34 upland cotton samples collected from Palo Verde Valley (1-24) and Coachella Valley (25-34) in California using double-antibody sandwich (DAS)-ELISA and RT-PCR assays specific for cotton leafroll dwarf virus.

| No.<sup>a</sup> | Symptoms<sup>b</sup> | ELISA | RT-PCR | No. | Symptoms | ELISA | RT-PCR |
| --- | --- | --- | --- | --- | --- | --- | --- |
| 1 | T + SI | - | - | 18 | T + SI | + | - |
| 2 | M | - | - | 19 | M | + | - |
| 3 | M | - | - | 20 | M | + | + |
| 4 | LR (aphids) | + | - | 21 | T + SI | - | - |
| 5 | LC | - | - | 22 | T + SI | + | - |
| 6 | LC | - | - | 23 | LC | - | - |
| 7 | M | + | - | 24 | T + SI | - | - |
| 8 | M | + | - | 25 | T + SI | + | - |
| 9 | LC and M | + | + | 26 | IC | + | - |
| 10 | M | - | - | 27 | IC | - | + |
| 11 | LC | - | - | 28 | IC | - | - |
| 12 | Br | - | - | 29 | IC | - | - |
| 13 | LC | - | - | 30 | IC | - | - |
| 14 | T + SI | - | - | 31 | IC | - | - |
| 15 | T + SI, LR | + | + | 32 | IC | - | + |
| 16 | LR | + | - | 33 | IC | + | - |
| 17 | T + SI | + | - | 34 | T + SI | + | - |

<sup>a</sup> Samples 1-4 originated from PV1 location (unknown cultivar), 5-8 from PV2 location (DPL2127), 9-13 from PV3 location (unknown cultivar), 14-17 from PV4 location (DPL2012), 18 from PV5 location (DPL2012), 19-20 from PV6 location (unknown cultivar), 21-24 from PV7 location (DPL 2020), and 25-34 from Coach location (unknown cultivar).
<sup>b</sup> Symptoms: T + SI = taller plants with terminal stacked nodes, M = mosaic, LR = leaf rolling (aphids = presence of aphids was noted in the plants), LC = leaf crinkling, Br = bronzing, IC = interveinal chlorosis.

**Figure 2.**
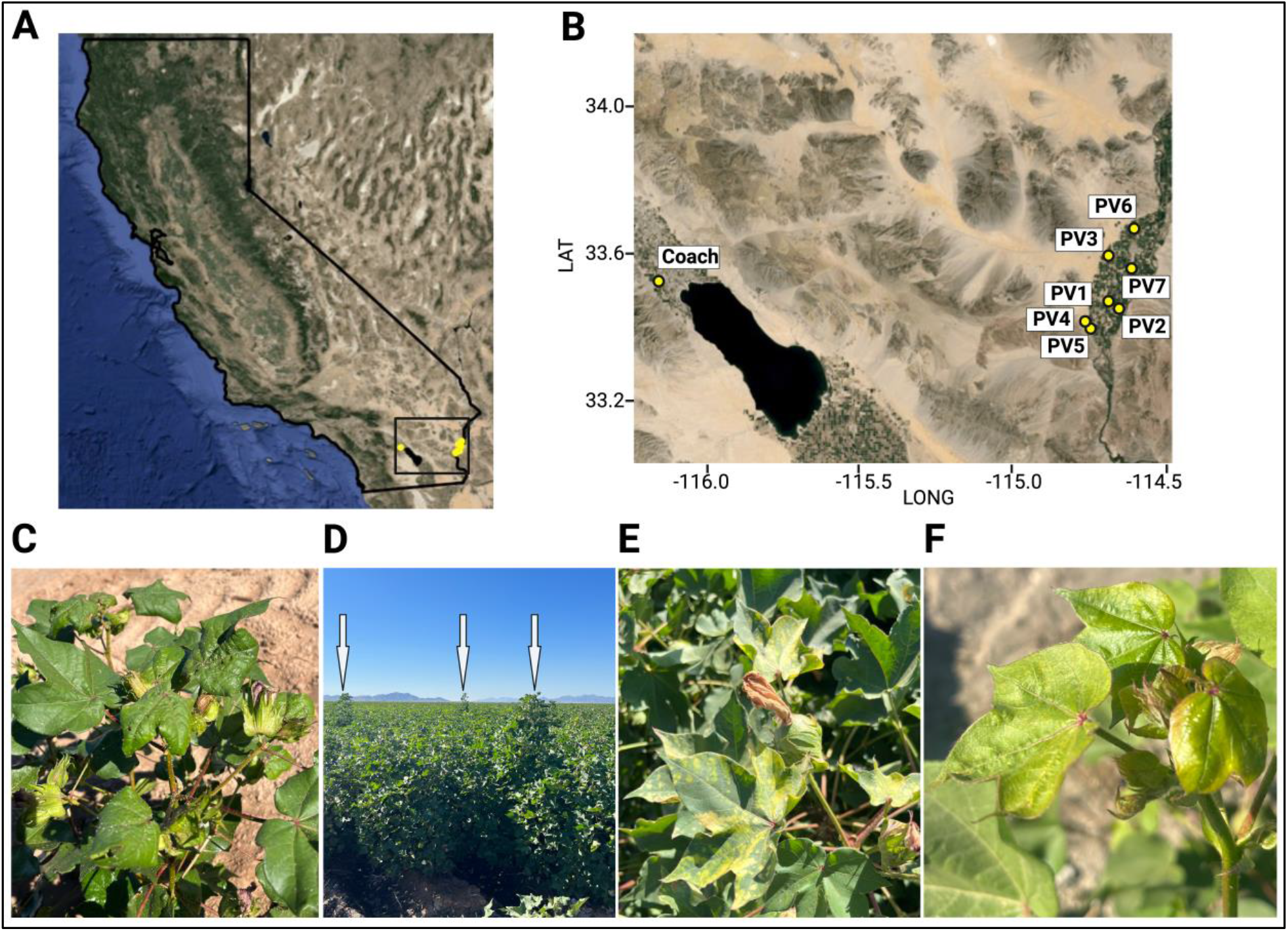
A) Locations in Southern California and zoom (B) where samples were collected from several fields across Palo Verde Valley (PV 1-7) and the Coachella Valley Agricultural Research Station (Coach). Symptoms: C) Leaf rolling and aphid presence (note abundance of white exoskeletons on leaves due to aphid molting), D) taller plants than their peers with stacked terminal nodes, and E) mosaic, observed in samples collected from Palo Verde Valley. Symptoms observed in the Coachella Valley Agricultural Research Station were characterized by leaf rolling and interveinal chlorosis (F).

**Figure 3.**
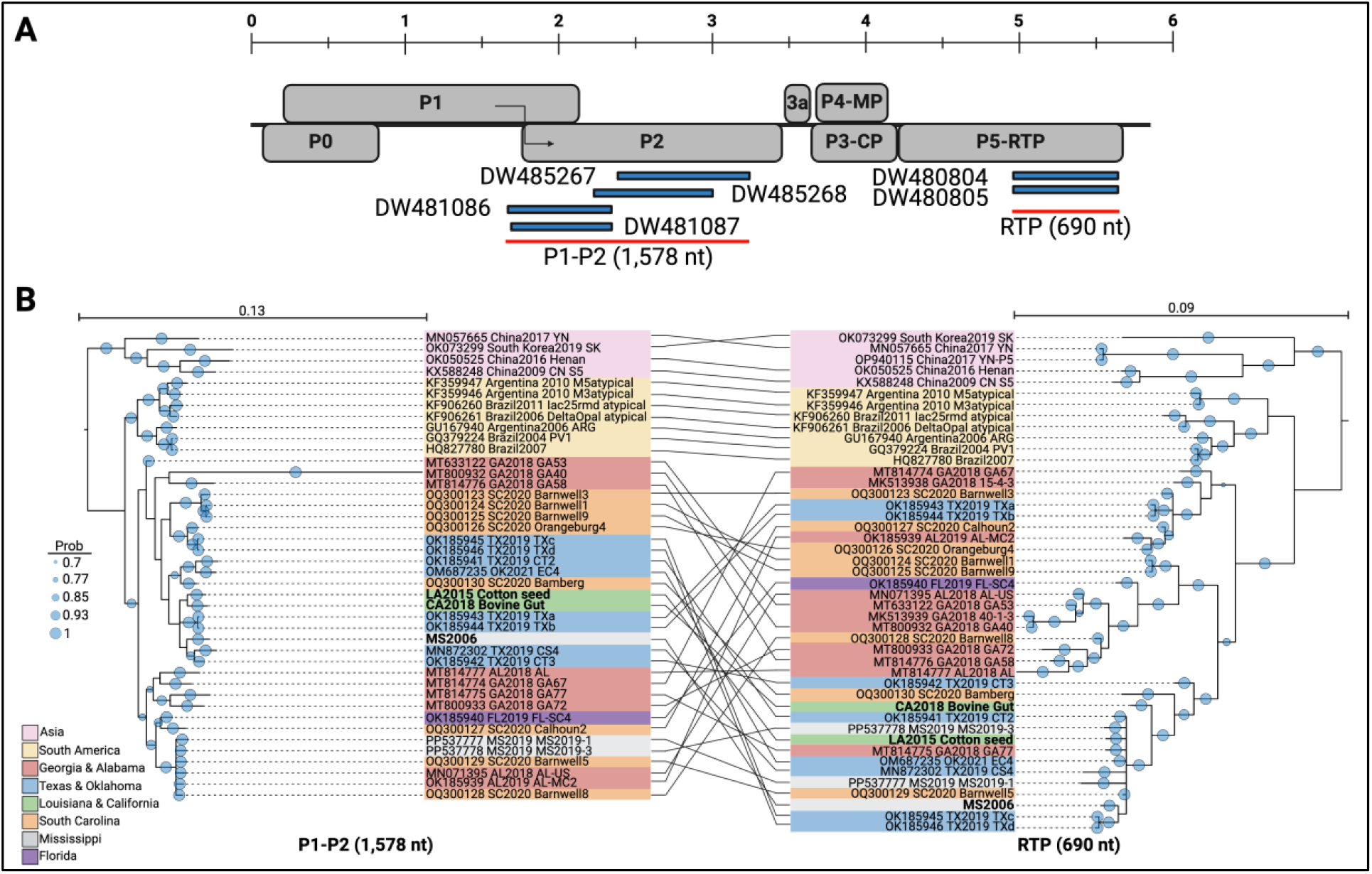
A) Representation of the expressed sequence tags (EST) *Gossypium hirsutum* accessions from 2006 that mapped to the cotton leafroll dwarf virus genome from Georgia (MT800932). Note: P1-P2 polyprotein is coded through a ribosomal frameshift that extends P1 towards P2 ORFs. Also, a CP-RTP protein is coded through a ribosomal readthrough extending P3 (coat protein, CP) towards P5 (read-through protein, RTP) ORFs. B) Phylogenetic trees constructed by Bayesian inference of consensus sequences obtained from the EST accessions mapping to the P1-P2 (1,578 nt)), and the RTP (690 nt) coding regions and their respective homologs in GenBank as of March 2024.

In the second data mining approach to retrieve additional CLRDV sequences, we built upon methodologies previously described by Bejerman and colleagues (Bejerman, Dietzgen, and Debat 2021) with some modifications. Briefly, the nucleotide sequences of the reference CLRDV isolate (NC014545), and a selected US isolate characterized from Georgia (MT800932) were used as queries in BLASTn searches against the Viridiplantae (taxid = 3193) transcriptome shotgun assembly (TSA), and the expressed sequence tags (est) databases using the following modified parameters: word size = 15 and expect threshold = 0.001. The EST database contains short ‘single-pass’ sequences of usually less than 1,000 bp that were generated from cDNA libraries cloned and sequenced using Sanger sequencing (Boguski, Lowe, and Tolstoshev 1993). The TSA database search yielded no significant hits (nucleotide identities <73.8% and query coverage <39%), suggesting the identified sequences were not closely related to CLRDV (data not shown). Conversely, the EST database search revealed six intriguing *G. hirsutum* accessions (DW480804-5, DW481086-7, and DW485267-8) with high nucleotide identities (87.5 – 96.8%) spanning the P1-P2 and P5 coding regions (Figure 3A). Interestingly, these EST sequences originated from Mississippi in 2006 and were derived from cDNA libraries constructed from RNA extracted from a variety of cotton tissues. These results strongly suggest the presence of CLRDV in the DES 119 upland cotton plants used by the researchers back in 2006. DES 119 was a popular cultivar in the southern US and was developed in the Delta Branch Mississippi Agricultural and Forestry Experiment Station in Stoneville, Mississippi (Bridge 1986).

Subsequently, we retrieved the EST accessions and generated consensus sequences for the P1-P2 (1,578 nt) and RTP (690 nt) coding regions. BLAST searches revealed high nucleotide identities: 98.6% for P1-P2 with a Texas isolate (OK185942) and 99.3% for RTP with a South Carolina isolate (OQ300129). To investigate evolutionary relationships, we constructed phylogenetic trees using the P1-P2 and RTP consensus sequences and their CLRDV homologs retrieved as detailed above. In this analysis, we also included the sequence homologs from two CLRDV isolates from Mississippi originally collected by our group in 2019 (GenBank accessions PP537777-8). The phylogenetic analyses of both P1-P2 and RTP (Figure 3B) presented congruent topology to the phylogenetic tree from almost complete genomes that presented geographically distinct clades (Figure 1B). Notably, the P1-P2 region from Mississippi 2006 clustered with CLRDV isolates from Texas, similar to the LA2015 and CA2018 isolates characterized in this study. Nevertheless, the RTP region from Mississippi 2006 and the LA2015 and CA2018 grouped with isolates from Texas, Georgia, and South Carolina (Figure 3B). These results further suggest the presence of CLRDV in Mississippi since at least 2006, likely undetected due to a possible lack of noticeable symptoms. CLRDD and CBD share a range of symptoms not only between each other but also with bronze wilt, a malady with unclear etiology that is characterized by reddening of leaves and stems, fruit abortion, leaf drooping, and wilting (Creech and Fieber 2000; Bell 2001). Bronze wilt is most severe in ‘short-season’ varieties derived from ‘Tamcot SP-37’ or ‘Miscot T8-27’ and emerged in Mississippi and Louisiana in 1995, but subsequently spread to other states: Texas, North Carolina, Tennessee, Georgia, Arkansas, and California (Bell 2001). It is plausible to hypothesize that CLRDV might induce bronze wilt symptoms in susceptible varieties in adequate environmental settings: high temperatures and dry conditions. Studies using infectious clones or controlled aphid-mediated transmission, and adequate environmental conditions in susceptible cultivars may validate this hypothesis.

Lastly, retrospective indexing of archived RNA or tissue samples from any state in the cotton belt where perhaps CLRDV was present before its official report in the country in 2017 could validate our computational findings. This will be of paramount importance to better understand the dynamics of the timeline and geographic spread of CLRDV in the US.

---

**Table S1.** Sequence Read Archive (SRA) accessions that Serratus (serratus.io) predicted to contain landmarks of the RNA-dependent RNA polymerase of cotton leafroll dwarf virus (CLRDV) and their specific metadata.

| No. | Accession | Organism | Country | Collection Date | No. reads | Virus Reads (%) | CLRDV reads (%) | CLRDV Reads (No.) |
| --- | --- | --- | --- | --- | --- | --- | --- | --- |
| 1 | SRR6379575 | *Gossypium hirsutum* | China | Unspecified | 51,631,110 | 0.006 | 0.00348 | 1797 |
| 2 | SRR6466408 | *G. hirsutum* | China | Unspecified | 56,294,020 | 0.03 | 0.0261 | 14693 |
| 3 | SRR6466409 | *G. hirsutum* | China | Unspecified | 56,031,882 | 0.2 | 0.192 | 107581 |
| 4 | SRR6466411 | *G. hirsutum* | China | Unspecified | 56,934,308 | 0.7 | 0.693 | 394555 |
| 5 | SRR6466412 | *G. hirsutum* | China | Unspecified | 56,513,454 | 0.3 | 0.291 | 164454 |
| 6 | SRR6466452 | *G. hirsutum* | China | Unspecified | 56,858,458 | 0.00004 | 0.000038 | 22 |
| 7 | SRR6466453 | *G. hirsutum* | China | Unspecified | 56,685,623 | 0.2 | 0.196 | 111104 |
| 8 | SRR6466454 | *G. hirsutum* | China | Unspecified | 55,641,875 | 0.1 | 0.095 | 52860 |
| 9 | SRR6466460 | *G. hirsutum* | China | Unspecified | 55,570,322 | 0.04 | 0.0376 | 20894 |
| 10 | SRR6466461 | *G. hirsutum* | China | Unspecified | 56,787,020 | 0.007 | 0.00028 | 159 |
| 11 | SRR6466462 | *G. hirsutum* | China | Unspecified | 55,251,426 | 0.02 | 0.0074 | 4089 |
| 12 | SRR6473051 | *G. hirsutum* | China | Unspecified | 56,294,015 | 0.03 | 0.0261 | 14693 |
| 13 | SRR6473052 | *G. hirsutum* | China | Unspecified | 56,031,890 | 0.2 | 0.192 | 107581 |
| 14 | SRR7820592 | *G. hirsutum* | China | Unspecified | 24,464,103 | 0.001 | 0.0005 | 122 |
| 15 | SRR8089834 | *G. hirsutum* | China | Unspecified | 27,671,165 | 0.01 | 0.0047 | 1301 |
| 16 | SRR8089835 | *G. hirsutum* | China | Unspecified | 32,087,658 | 0.009 | 0.00459 | 1473 |
| 17 | SRR8089836 | *G. hirsutum* | China | Unspecified | 25,902,332 | 0.02 | 0.0124 | 3212 |
| 18 | SRR8089837 | *G. hirsutum* | China | Unspecified | 27,913,778 | 0.01 | 0.0049 | 1368 |
| 19 | SRR8089838 | *G. hirsutum* | China | Unspecified | 30,052,416 | 0.04 | 0.0224 | 6732 |
| 20 | SRR8089839 | *G. hirsutum* | China | Unspecified | 31,885,100 | 0.05 | 0.0375 | 11957 |
| 21 | SRR8089840 | *G. hirsutum* | China | Unspecified | 23,956,823 | 0.0007 | 0.000343 | 82 |
| 22 | SRR8089843 | *G. barbadense* | China | Unspecified | 33,195,430 | 0.05 | 0.033 | 10954 |
| 23 | SRR8089857 | *G. barbadense* | China | Unspecified | 22,139,250 | 0.05 | 0.032 | 7085 |
| 24 | SRR8089865 | *G. hirsutum* | China | Unspecified | 22,920,974 | 0.0005 | 0.000305 | 70 |
| 25 | SRR8089867 | *G. hirsutum* | China | Unspecified | 21,829,353 | 0.004 | 0.00176 | 384 |
| 26 | SRR8089868 | *G. hirsutum* | China | Unspecified | 25,521,887 | 0.01 | 0.0075 | 1914 |
| 27 | SRR8089870 | *G. hirsutum* | China | Unspecified | 22,360,056 | 0.04 | 0.0292 | 6529 |
| 28 | SRR8089872 | *G. barbadense* | China | Unspecified | 22,568,216 | 0.006 | 0.0033 | 745 |
| 29 | SRR8089873 | *G. barbadense* | China | Unspecified | 22,360,056 | 0.04 | 0.04 | 8944 |
| 30 | SRR8089884 | *G. barbadense* | China | Unspecified | 27,617,866 | 0.007 | 0.007 | 1933 |
| 31 | SRR8089889 | *G. barbadense* | China | Unspecified | 28,710,754 | 0.0003 | 0.0003 | 86 |
| 32 | SRR8089902 | *G. hirsutum* | China | Unspecified | 23,712,961 | 0.005 | 0.00205 | 486 |
| 33 | SRR8089905 | *G. barbadense* | China | Unspecified | 22,161,291 | 0.0003 | 0.000231 | 51 |
| 34 | SRR8089933 | *G. barbadense* | China | Unspecified | 27,245,807 | 0.00007 | 0.0000392 | 11 |
| 35 | SRR8089937 | *G. barbadense* | China | Unspecified | 24,677,326 | 0.006 | 0.00408 | 1007 |
| 36 | SRR8089938 | *G. barbadense* | China | Unspecified | 26,412,978 | 0.002 | 0.00138 | 364 |
| 37 | SRR8089939 | *G. barbadense* | China | Unspecified | 25,205,732 | 0.02 | 0.0144 | 3630 |
| 38 | SRR8089940 | *G. barbadense* | China | Unspecified | 24,220,501 | 0.03 | 0.0231 | 5595 |
| 39 | SRR8089941 | *G. barbadense* | China | Unspecified | 26,972,409 | 0.01 | 0.0075 | 2023 |
| 40 | SRR8089942 | *G. barbadense* | China | Unspecified | 26,972,409 | 0.01 | 0.0075 | 2023 |
| 41 | SRR8089943 | *G. barbadense* | China | Unspecified | 35,349,902 | 0.02 | 0.0146 | 5161 |
| 42 | SRR8089944 | *G. barbadense* | China | Unspecified | 24,258,917 | 0.0004 | 0.000268 | 65 |
| 43 | SRR8089945 | *G. barbadense* | China | Unspecified | 21,470,213 | 0.004 | 0.004 | 859 |
| 44 | SRR8089946 | *G. barbadense* | China | Unspecified | 33,843,907 | 0.01 | 0.01 | 3384 |
| 45 | SRR8089947 | *G. barbadense* | China | Unspecified | 24,838,024 | 0.001 | 0.001 | 248 |
| 46 | SRR8090013 | *G. barbadense* | China | Unspecified | 22,782,729 | 0.002 | 0.002 | 456 |
| 47 | SRR8090043 | *G. hirsutum* | China | Unspecified | 26,665,033 | 0.0001 | 0.0001 | 27 |
| 48 | SRR8090051 | *G. barbadense* | China | Unspecified | 26,256,317 | 0.1 | 0.1 | 26256 |
| 49 | SRR8090079 | *G. barbadense* | China | Unspecified | 28,717,540 | 0.005 | 0.005 | 1436 |
| 50 | SRR10336136 | *G. hirsutum* | China | 8/30/15 | 47,400,992 | 0.07 | 0.0693 | 32849 |
| 51 | SRR10336137 | *G. hirsutum* | China | 8/30/15 | 52,804,231 | 0.02 | 0.019 | 10033 |
| 52 | SRR10336138 | *G. hirsutum* | China | 8/30/15 | 47,319,631 | 0.009 | 0.00873 | 4131 |
| 53 | SRR10336139 | *G. hirsutum* | China | 8/30/15 | 54,202,766 | 0.02 | 0.019 | 10299 |
| 54 | SRR10336140 | *G. hirsutum* | China | 7/30/15 | 55,210,877 | 0.004 | 0.00344 | 1899 |
| 55 | SRR10336143 | *G. hirsutum* | China | 7/30/15 | 51,650,145 | 0.005 | 0.00495 | 2557 |
| 56 | SRR10336144 | *G. hirsutum* | China | 8/30/15 | 61,114,750 | 0.02 | 0.0196 | 11978 |
| 57 | SRR10336145 | *G. hirsutum* | China | 8/30/15 | 55,527,415 | 0.03 | 0.0294 | 16325 |
| 58 | SRR10336146 | *G. hirsutum* | China | 8/30/15 | 52,730,844 | 0.02 | 0.0196 | 10335 |
| 59 | SRR10336147 | *G. hirsutum* | China | 8/30/15 | 43,389,500 | 0.02 | 0.0196 | 8504 |
| 60 | SRR10336148 | *G. hirsutum* | China | 8/30/15 | 50,611,246 | 0.05 | 0.0495 | 25053 |
| 61 | SRR10336149 | *G. hirsutum* | China | 7/30/15 | 55,825,073 | 0.003 | 0.00297 | 1658 |
| 62 | SRR10336150 | *G. hirsutum* | China | 7/30/15 | 46,163,542 | 0.03 | 0.0297 | 13711 |
| 63 | SRR10914150 | *Hibiscus cannabinus* | China | Unspecified | 26,311,274 | 0.003 | 0.00213 | 560 |
| 64 | SRR11191655 | *G. hirsutum* | China | Unspecified | 37,488,038 | 0.01 | 0.0076 | 2849 |
| 65 | SRR11191718 | *G. hirsutum* | China | Unspecified | 36,436,476 | 0.04 | 0.0392 | 14283 |
| 66 | SRR12316179 | *G. hirsutum*<br>*x G. australe* | China | Unspecified | 30,551,723 | 0.002 | 0.00198 | 605 |
| 67 | SRR12316181 | *G. hirsutum*<br>*x G. australe* | China | Unspecified | 38,056,129 | 0.003 | 0.003 | 1142 |
| 68 | SRR14374199 | *G. barbadense* | China | Unspecified | 59,420,294 | 0.003 | 0.00246 | 1462 |
| 69 | SRR14374200 | *G. barbadense* | China | Unspecified | 72,841,338 | 0.02 | 0.0172 | 12529 |
| 70 | SRR14374201 | *G. barbadense* | China | Unspecified | 78,634,409 | 0.2 | 0.186 | 146260 |
| 71 | SRR14826282 | *G. hirsutum* | China | Unspecified | 19,281,435 | 0.02 | 0.018 | 3471 |
| 72 | SRR14826285 | *G. hirsutum* | China | Unspecified | 23,014,574 | 0.007 | 0.00392 | 902 |
| 73 | SRR15495736 | *G. hirsutum* | China | Unspecified | 33,275,652 | 0.005 | 0.00245 | 815 |
| 74 | SRR1562253 | *G. hirsutum* | China | Unspecified | 54,430,237 | 0.007 | 0.00672 | 365771 |
| 75 | SRR1695179 | *G. hirsutum* | China | Unspecified | 12,331,675 | 0.002 | 0.002 | 24663 |
| 76 | SRR18105835 | *G. hirsutum* | China | Unspecified | 28,063,971 | 0.0006 | 0.0000048 | 135 |
| 77 | SRR18105836 | *G. barbadense* | China | Unspecified | 28,228,236 | 0.001 | 0.0000082 | 231 |
| 78 | SRR18105837 | *G. barbadense* | China | Unspecified | 27,654,865 | 0.001 | 0.000008 | 221 |
| 79 | SRR18907716 | *G. hirsutum* | China | Unspecified | 31,159,241 | NA | NA | NA |
| 80 | SRR18907718 | *G. hirsutum* | China | Unspecified | 26,403,948 | NA | NA | NA |
| 81 | SRR18907719 | *G. hirsutum* | China | Unspecified | 25,336,916 | NA | NA | NA |
| 82 | SRR6012273 | *G. hirsutum* | China | Unspecified | 231,287,452 | NA | NA | NA |
| 83 | SRR6012493 | *G. hirsutum* | China | Unspecified | 44,376,979 | 0.02 | 0.00005 | 2219 |
| 84 | SRR8289730 | *G. hirsutum* | China | Unspecified | 22,759,247 | 0.05 | 0.00049 | 11152 |
| 85 | SRR8289731 | *G. hirsutum* | China | Unspecified | 21,468,820 | 0.02 | 0.000188 | 4036 |
| 86 | SRR8289732 | *G. hirsutum* | China | Unspecified | 30,175,804 | 0.009 | 0.0000891 | 2689 |
| 87 | SRR8289733 | *G. hirsutum* | China | Unspecified | 35,523,909 | 0.006 | 0.000054 | 1918 |
| 88 | SRR8289734 | *G. hirsutum* | China | Unspecified | 25,947,904 | 0.0007 | 0.00000686 | 178 |
| 89 | SRR8289735 | *G. hirsutum* | China | Unspecified | 28,153,497 | 0.02 | 0.000196 | 5518 |
| 90 | SRR8289736 | *G. hirsutum* | China | Unspecified | 23,734,295 | 0.003 | 0.0000219 | 520 |
| 91 | SRR8289737 | *G. hirsutum* | China | Unspecified | 22,371,665 | 0.003 | 0.0000204 | 456 |
| 92 | SRR8289740 | *G. hirsutum* | China | Unspecified | 21,270,820 | 0.002 | 0.0000064 | 136 |
| 93 | SRR8289741 | *G. hirsutum* | China | Unspecified | 20,968,129 | 0.001 | 0.0000095 | 199 |
| 94 | SRR8289743 | *G. hirsutum* | China | Unspecified | 28,994,664 | 0.0007 | 0.0007 | 20296 |
| 95 | SRR8289744 | *G. hirsutum* | China | Unspecified | 26,549,541 | 0.001 | 0.001 | 26550 |
| 96 | SRR8164488 | *G. hirsutum* | India | Unspecified | 26,216,384 | 0.002 | 0.00186 | 48762 |
| 97 | SRR14368410 | *G. hirsutum* | India | Unspecified | 51,011,713 | 0.3 | 0.003 | 153035 |
| 98 | SRR14368411 | *G. hirsutum* | India | Unspecified | 45,289,016 | 0.3 | 0.003 | 135867 |
| 99 | SRR14368412 | *G. hirsutum* | India | Unspecified | 44,377,225 | 0.3 | 0.003 | 133132 |
| 100 | SRR14368413 | *G. hirsutum* | India | Unspecified | 39,613,701 | 0.36 | 0.00288 | 114087 |
| 101 | SRR14368414 | *G. hirsutum* | India | Unspecified | 57,259,502 | NA | NA | NA |
| 102 | SRR14368415 | *G. hirsutum* | India | Unspecified | 55,167,588 | 0.06 | 0.000006 | 331 |
| 103 | SRR14368416 | *G. hirsutum* | India | Unspecified | 43,791,288 | 0.05 | 0.000005 | 219 |
| 104 | SRR14368417 | *G. hirsutum* | India | Unspecified | 38,307,956 | 0.05 | 0.000005 | 192 |
| 105 | SRR14368418 | *G. hirsutum* | India | Unspecified | 56,146,907 | 0.3 | 0.00003 | 1684 |
| 106 | SRR14368419 | *G. hirsutum* | India | Unspecified | 51,040,836 | 0.3 | 0.00003 | 1531 |
| 107 | SRR14368420 | *G. hirsutum* | India | Unspecified | 39,618,132 | NA | NA | NA |
| 108 | SRR14368421 | *G. hirsutum* | India | Unspecified | 40,160,195 | 0.05 | 0.00002 | 803 |
| 109 | SRR16573720 | *Linum*<br>*usitatissimum* | India | Unspecified | 24,253,258 | 0.004 | 0.000032 | 776 |
| 110 | SRR22177514 | *Cicer arietinum* | India | Unspecified | 28,735,806 | 0.01 | 0.00003 | 862 |
| 111 | SRR8164489 | *G. hirsutum* | India | Unspecified | 36,731,948 | 0.004 | 0.004 | 146928 |
| 112 | SRR10765499 | *G. hirsutum* | USA | 8/15/19 | 21,877,261 | 0.02 | 0.0126 | 275653 |
| 113 | SRR1805340 | *G. hirsutum* | USA | Unspecified | 82,059,811 | 0.02 | 0.0194 | 1591960 |
| 114 | SRR9035366 | Bovine Gut<br>Metagenome | USA | 5/13/18 | 39,269,410 | 0.41 | 0.0002 | 7854 |

NA = information not available for that specific dataset.

